# Data leakage and measurement error inflate the apparent predictability of overyielding from plant traits

**DOI:** 10.64898/2026.01.15.699684

**Authors:** Emanuel B. Kopp, Noelle Koenig, Lukas Vonmetz, Samuel E. Wuest, Pascal A. Niklaus

## Abstract

Predicting plant mixture overyielding from functional traits is central to understanding how biodiversity influences ecosystem functioning. Combining empirical data from a large mixture experiment (764 mixtures) and two trait-measurement experiments containing 90 soybean genotypes, with complementary simulations where trait-function relationships and noise levels were defined a priori, we show that apparent model performance depends critically on how predictive models are validated. When training and testing data share genotypes, predictive ability is strongly inflated because shared monoculture and trait measurements create data leakage. Measurement errors further propagate through these shared components, inducing spurious correlations and amplifying noise. When validation uses completely independent genotypes, the predictive power of both linear and machine-learning models declines sharply, revealing limited but genuine predictability. These results show how data structure and measurement error can produce misleading model performance and underscore the need for rigorous validation to achieve robust ecological prediction.

## Introduction

It is well established that more species-rich plant communities are on average more productive than monocultures, a phenomenon termed overyielding (Hector et al. 1999; Hooper et al. 2005; Cardinale et al. 2007; Huang et al. 2018). In agriculture, variety mixtures, i.e. the mixed cultivation of different varieties of the same crop, promise higher yields and could easily be integrated into modern production schemes (Wuest et al. 2021; Kopp et al. 2023; Huang et al. 2024). However, variety mixtures differ greatly with respect to overyielding (Reiss and Drinkwater 2018), and it is currently very difficult to predict which mixtures overyield and to which extent (Borg et al. 2018). This limits the development and practical use of variety mixtures in agriculture. It therefore is urgent to develop tools to effectively predict variety mixtures that promise overyielding (Kopp et al. 2023).

Ecological theory suggests that the overyielding of mixtures of species or varieties results from niche complementarity among species or varieties (Godoy et al. 2020; Amyntas et al. 2023). However, niche complementarity is a broad concept encompassing different mechanisms such as resource partitioning, abiotic facilitation, and biotic feedback (Barry et al. 2019). Traditionally, niches are described as a set of environmental conditions allowing a species’ population to maintain a positive growth rate (Chase and Leibold 2003). Geometrically, niches have been conceptualized as hypervolumes where the dimensions are environmental conditions (Hutchinson 1957). However, niche hypervolumes may have complex niche geometries, including concave shapes and holes, which are complicated to describe (Blonder 2018). It also is unclear what dimensions of niche space are, and the niche concept thus remains elusive and niches difficult to quantify. Instead of using environmental conditions, ecologists often use functional traits to describe niches. Under this premise, distinct functional traits may indicate functional niche complementarity (Ebeling et al. 2014; Wagg et al. 2017; Mahaut et al. 2023). However, the relationship between niche-space and trait-space is not straightforward (e.g.Violle and Jiang 2009), and niches may geometrically be as complicated, or even more complicated, when analysed in trait space than when using environmental dimensions as axes of niche space.

Quantifying how niches translate into ecosystem outcomes, such as overyielding, typically requires predictive models based on species or variety traits. To predict overyielding based on the traits of the varieties that are co-cultivated, different types of models may be used. To compare the performance of these models, one option are information criteria (e.g. AIC or BIC), which balance the fit of a model with its complexity (e.g. the number of parameters estimated, see Yates et al. 2023). Another option is systematic cross-validation, in which datasets are split into “training data” to estimate model parameter, and “test data” for the evaluation of the model’s predictive ability on independent data (Roberts et al. 2017). When cross-validating, it is very important to avoid data leakage from the training into the test dataset. Data leakage has been described as the “illicit sharing of information between the training data and the test data” (Bernett et al. 2024). Under leakage, the predictive performance of a model often depends on data that is not causally related to the predicted pattern, but that indirectly describes some association between training and test data. This may, for example, occur when the training and test data overlap, and the common items in training and test dataset are identified using attributes that are not causally related to the phenomenon that is modeled. When the model is applied to completely independent data, this shared information is no longer present and the predictive ability collapses.

As part of a larger research endeavor, we have been studying overyielding in soybean variety mixtures. Here, we are concerned with a series of traits that we measured in 90 soybean varieties and which we used to build models to predict the overyielding of binary mixtures of these varieties. In a first step, we tested different types of predictive models and evaluated their performance using different cross-validation schemes. A challenge with such an empirical comparison of model performances is that the true underlying quantitative relationship between traits and overyielding remains unknown. To more rigorously evaluate advantages and disadvantages of our models and cross-validation schemes, we therefore repeated the same analyses and cross-validation, this time using data simulated with known trait-effect relationships. Comparing all these data and approaches, we provide practical guidance for predicting mixture overyielding from traits while avoiding common pitfalls like data leakage and model overfitting.

## Material and methods

### Empirical data collection and trait processing

Ninety soybean (*Glycine max*) varieties were selected based on their geographic origin and domestication history (ranging from landraces to elite cultivars, all early-maturing to suit the growth conditions of Switzerland and most obtained from the USDA germplasm collection – see supplementary materials for details). For all experiments, seeds were inoculated with rhizobia (HiStick, BASF, containing *Bradyrhizobium japonicum*) and germinated in propagator trays in the greenhouse. Variety mixture data was obtained from a pot experiment (Fig. 1A, hereafter called mixture experiment) conducted during the summer 2022 in Wädenswil, Switzerland (47.222°N, 8.669°E, 509 m a.s.l.). After 14 days in the propagator trays, the seedlings were transplanted pairwise into 5 l pots and inoculated with 10 ml of a solution of 10 g/l of mycorrhizal suspension (*Rhizophagus irregularis*, Lalrise Max WP, Lallemand Inc, Canada). From the 90 varieties used, 8 varieties differing in geographic origin and domestication history were selected as so-called “testers”. All 90 varieties were paired with these 8 testers for a total of 712 distinct two-way mixtures and 8 tester monocultures. All variety monocultures were replicated twice. The pots were arranged into two blocks. Each block further contained 41 distinct two-way mixtures from the 82 non-tester varieties to increase the total number of mixtures. Hence, the final experiment consisted of 2 blocks with 712 mixtures containing tester varieties, 41 mixtures not containing tester varieties, 24 tester monocultures and 164 non-tester monocultures each, for a total of 1882 pots or 3764 plants (Fig. 1B). The monocultures and mixtures were grown under hail protection nets and irrigated daily. Each variety was individually harvested after reaching maturity. All harvested plants were dried for at least 24h at 60°C and shoot biomass weighed.

**Figure 1:**
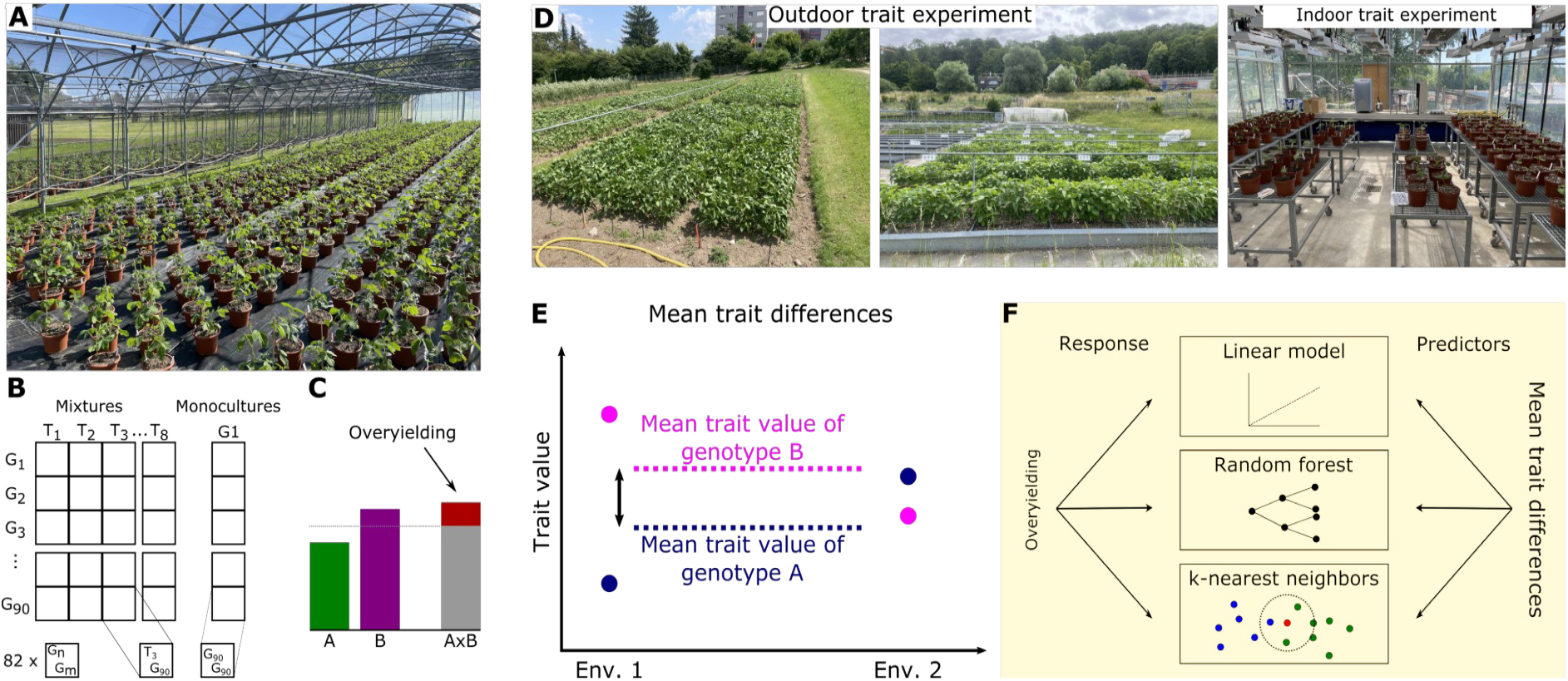
(A) the overyielding experiment, in each pot two plant of either the same variety (monoculture) or different varieties (mixture) grow together. (B) Experimental design. (C) Overyielding representes the deviation of the observed mixture productivity from the expectation based on the mixture components’ productivity as monocultures. (D) The trait experiment. (E) Calculation of mean trait differences for a pair of genotypes. (F) Modelling approach: mean trait differences are used as predictors in three different model types to predict overyielding.

Genotype functional traits, which we used as predictors of mixture overyielding, were obtained from two experiments conducted in 2022 and in 2023 (hereafter called trait experiments; Fig. 1D). In the first experiment, the 90 soybean varieties were grown as single plants in 5 l pots arranged in four blocks in the greenhouse (total of 360 pots). In the second trait experiment, the same 90 varieties were grown outdoors at two different locations (Wädenswil, 47.222°N, 8.669°E, 509 m a.s.l. and Zürich-Irchel, 47.396°N, 8.551°E, 506 m a.s.l; distance ca. 21 km). We grew the varieties in rows, with four plant individuals spaced 4 cm apart, leaving a 16 cm gap between groups of different varieties. Each variety was replicated in one to three such groups, depending on seed availability. The position of each variety was randomized, with the condition that replicates never had the same neighbors.

In both experiments, we measured final plant height, aboveground biomass, and seed yield. In the first experiment, we collected additional trait data including plant height growth rate, leaf number, size, and weight, chlorophyll content, stomatal conductance, and flowering date.

### Data processing

Data from the mixture trial was corrected for spatial effects, and overyielding of each mixture calculated as difference between mixture yield and the average of the monoculture yields of the two mixture components (Fig. 1C). Trait data were averaged by genotype, and trait differences calculated for all pairwise genotype combinations. (Fig. 1E).

### Modelling of Overyielding

We modelled the dependency of overyielding (OY) on traits using (1) linear models, (2) random forests, and (3) *k* -nearest-neighbor (kNN) models (Fixt and Hodges 1989; Ho 1995; Fig. 1F). To build the linear models, we used the lm function from GLM.jl (Bates et al. 2023) in Julia (v1.9.1). We started by including all variables as predictors and proceeded with backwards selection, i.e. by removing individual variables sequentially as long as the model AIC decreased, to obtain the final model. Random forest models were implemented through DecisionTrees.jl (Sadeghi et al. 2022) within the MLJ.jl framework (Blaom et al. 2024), using the RandomForestRegressor function with default values except for the number of trees, set to 500. kNN models were implemented through NearestNeighbors.jl (Carlsson et al. 2024) within the MLJ.jl framework using the kNNRegressor function with default parameter settings. kNN models simply place the predictor data in an Euclidean space (Fixt and Hodges 1989) and make predictions by averaging the values of the *k* nearest data points found in this space; in our study, we used k=3. For all models, we used the trait differences among genotypes as predictors.

### Cross-validation

We cross-validated our models in two ways (Fig. 2): in the first, which we refer to as “standard”, the test data consisted of all 82 mixtures not containing any tester genotype plus 2 additional randomly selected mixtures for each tester variety. In the second, which we refer to as “strict”, we randomly chose one of the eight tester varieties and 19 of the 82 non-tester varieties. The test data then consisted of all the mixtures assembled from these 20 varieties, while the training set consisted of all the mixtures composed from the remaining 70 varieties.

**Figure 2:**
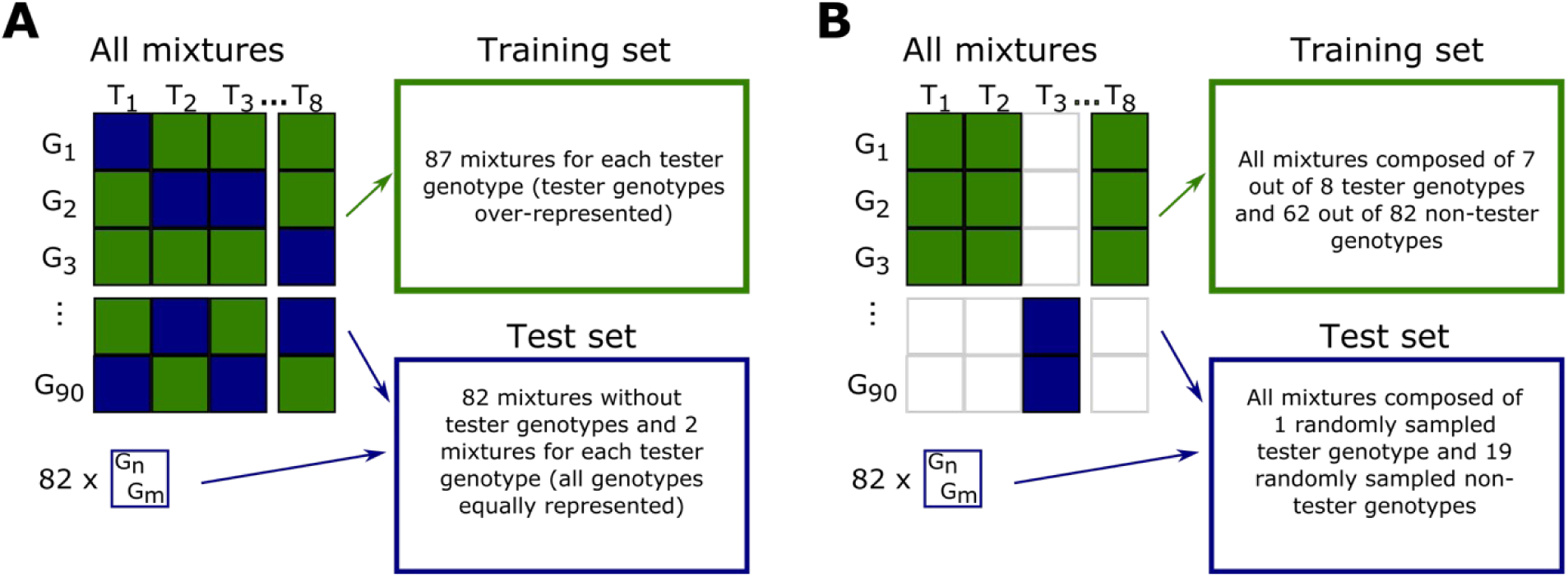
(A) Standard cross-validation: training set and test set do not share any specific mixtures, but have mixtures which share single components. (B) Strict cross-validation: training set and test set don’t share any genotypes.

The standard cross validation allowed us to use all data, achieving an 85:15 size ratio between training and test datasets. The random addition of two mixtures per tester genotype to the test dataset allowed us to run each model multiple times with slightly different training and test datasets to assess the stability of the predictive model. Overall, training and test datasets contained entirely different mixtures, while sharing some mixture components. The strict cross validation used only part of the data but had the advantage to avoid any overlap in the variety sets on which the mixtures of training and test datasets were based: not only were no mixture compositions shared, but the mixtures in training and test datasets were based on entirely different components.

Each model was run 50 times with different splits of the data into training and test subsets. For each run, the model was fit on the training data, and the expected OY of the test data was predicted. Predictive ability was quantified as the out-of-sample R^2^, calculated as the correlation between predicted and observed OY in the test set. To summarize performance across runs, we report the median out-of-sample R^2^, which is robust to extreme values, along with the interquartile range (25^th^ - 75^th^ percentiles) to indicate variability in predictive performance.

## Results

### Overyielding

Over the 764 mixtures analyzed (30 were lost because plants died or looked bad), shoot biomass overyielded on average by 4.4 g per pot (+3.5%). However, overyielding was highly variable, with estimated effects ranging from −51 to 94 g per pot (-35% to 72%, Fig. 3A). While negative overyielding is rare in natural communities (Cardinale et al. 2007), it does occur in crops and our findings are in line with corresponding studies (see Reiss and Drinkwater 2018 for a meta-analysis). Moreover, our OY estimates carry the compound biomass measurement error of the mixture biomass and the biomass of the two reference monocultures, and some of the observed variation in OY may thus be due to random measurement error.

**Figure 3:**
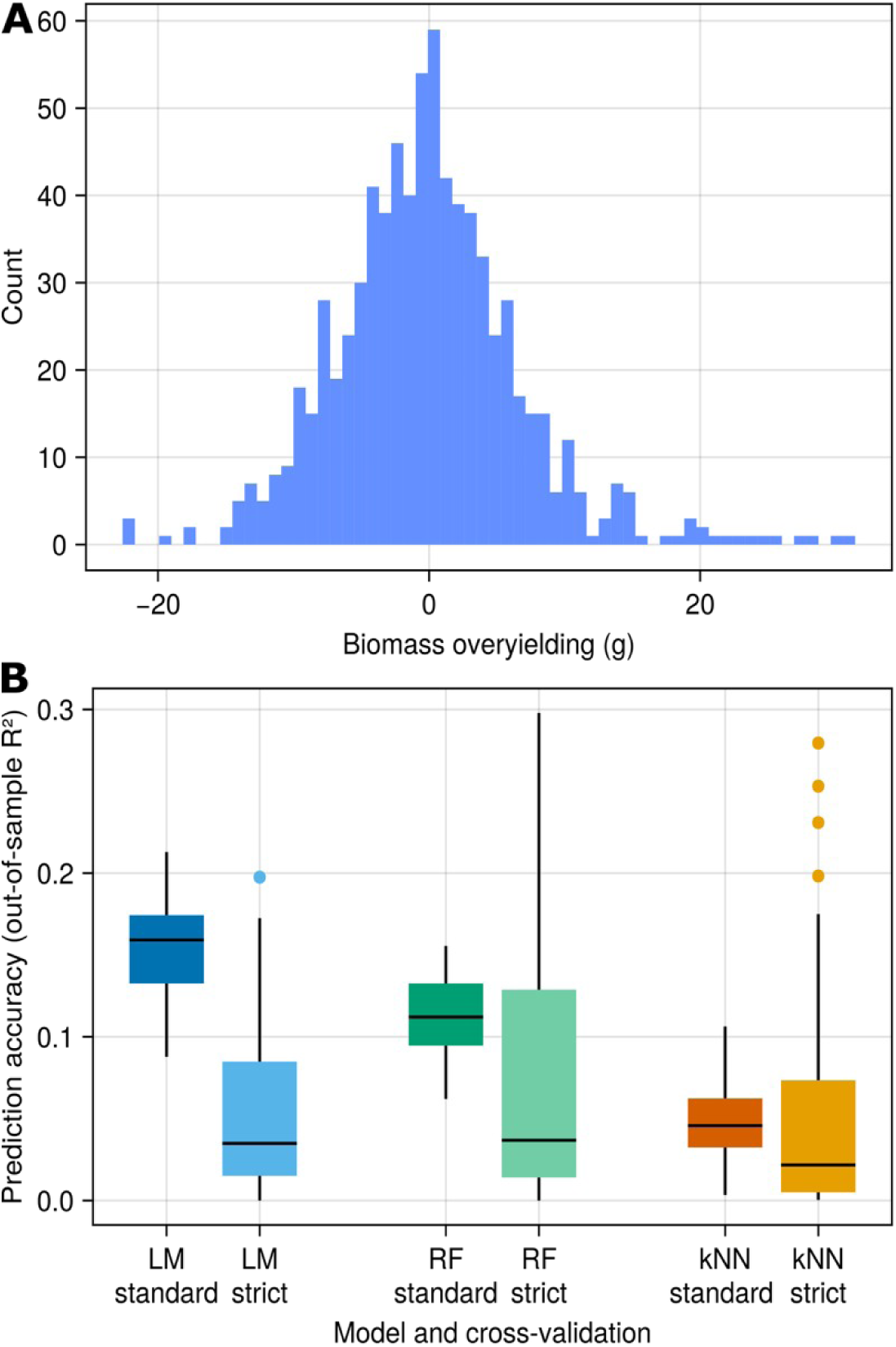
Variance in overyielding in experimental data (A) and prediction accuracy (out-of-sample R2) for the three tested models (LM: linear model; RF: random forest, kNN: k-nearest-neighbors) using standard or strict cross-validation.

Depending on the specific split between training and test data, the best linear model after backward variable selection contained between 5 and 12 predictors. Some predictors, for example the genotypic differences in produced biomass, seed yield, leaf dry matter content and time of flowering where present in all models, suggesting that they are correlated with plant-plant interactions that underlie overyielding. Other predictors, such as genotypic differences in leaf conductivity, specific leaf area, number of branches, or relative chlorophyll content, were never included in the best model.

### Cross-validation

With standard cross-validation, models explained more variance in overyielding observed in the test dataset than when using strict cross-validation. Linear models and random forest dropped substantially in predictive ability from standard to strict cross-validation, with out-of-sample R^2^, for LM, dropping from 0.159 (0.133 – 0.174) to 0.039 (0.017 – 0.104), and for random forest from 0.112 (0.095 – 0.133) to 0.037 (0.016 – 0.179). kNN showed limited predictive ability, with an out-of-sample R^2^ of 0.046 (0.032 – 0.062) under standard cross-validation and of 0.022 (0.005 – 0.093) under strict cross-validation.

The variability of the out-of-sample R^2^ generally was higher when using strict cross-validation than when using standard cross-validation (Fig. 3B). This is because in standard cross-validation, all 82 non-tester mixtures are always included in the test dataset, leading to higher similarity between the test sets among model runs. In contrast, with strict cross-validation, the test datasets were more variable among model runs. Moreover, in strict cross-validation both training and test dataset were smaller than in standard cross-validation.

With respect to data leakage, a key difference between the two cross-validation types is that in standard cross-validation, mixtures in the training and the test set share some components, whereas in the strict cross-validation, the mixtures in the two sets are assembled from entirely different varieties. One possibility therefore is that with standard cross-validation, the models “learn” to identify the presence of some varieties using trait combinations and then associates these varieties with specific overyielding effects. This is not possible with strict cross-validation. The same problem could also occur due to yield measurement error, since all overyielding estimates of mixtures involving a particular variety are based on the monoculture yield of this variety, i.e. a measurement error in monoculture yield will induce a positive correlation between overyielding estimates in training and test data, which the predictive model may pick up.

In summary, the reduced predictive performance of the models under strict cross-validation may have different origins, which are hard to separate with the present data. This includes noise (measurement error) that is shared between training and test datasets (monoculture values of shared varieties), or differences in the size and variability of training and test datasets. To investigate this issue more systematically, we therefore simulated datasets with known linear and non-linear relationships between genotypic trait differences and overyielding, adding varying levels of random noise.

### Simulation study

#### Study design and data simulation

We designed the simulation study to reflect the main features of our empirical study. Specifically, we simulated 30 genotypes and all possible pairwise combination of these (435 mixtures). For each genotype, we simulated 12 hypothetical plant functional traits, randomly sampled from a standard normal distribution, and calculated the trait differences between genotypes.

Mixture yields were simulated as the average yield of the components plus an overyielding component that depended on trait differences between varieties. We considered two functional forms for OY: a linear function of four selected trait differences, and a non-linear function of three trait differences. These functions serve as examples of possible *trait differences-OY relationships*, while the remaining traits varied independently and did not contribute to OY.

Observed yields included additive measurement error:

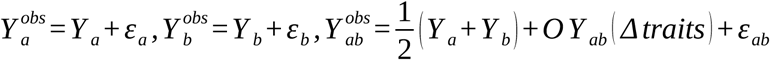

Observed overyielding was then calculated as the difference between observed mixture yield and the average of observed monocultures:

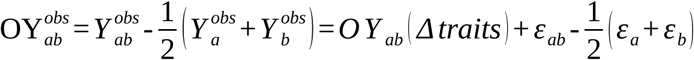

Measurement error in traits or yields can propagate into OY. Trait errors can create spurious correlations between traits and OY, while yield errors induce correlations among mixtures sharing a genotype, potentially generating data leakage between training and test sets. We evaluated model performance using two cross-validation schemes. In standard cross-validation, mixtures were randomly partitioned into training and test sets, allowing genotypes to occur in both datasets. In strict cross-validation, no genotype appeared in both training and test sets.

We repeated these simulation experiment 50 times for each cross-validation strategy. Additionally, we varied the noise level (standard deviation of the error term in the overyielding models) across a range of values to examine how increasing noise impacts model predictive performance. As we modelled the response explicitly, this allowed us to compute the signal-to-noise ratio (SNR) by dividing the variance of OY modelled without addition of noise by the variance of the OY noise. Again, we included linear regression with previous AIC-based backwards variable selection (LM), random forest (RF), and k-nearest neighbors (kNN). We performed all simulations using R (v4.3.1).

#### Simulation results

As expected, predictive performance decreased as noise increased (SNR decreased), regardless of model or cross-validation type (Fig 4). Linear models performed well under linear trait-OY relationships, particularly at higher SNRs, getting close to the theoretical maximum (the R^2^ explained by a model using the known relationship used to model yield; Fig. 4A). Random forest showed lower predictive performance compared to LM under standard cross-validation, and the performance dropped significantly under strict cross-validation, suggesting that much of its apparent power in standard cross-validation was due to leakage. kNN was consistently outperformed by the other models and showed a strong decline in predictive power under strict cross-validation.

**Figure 4:**
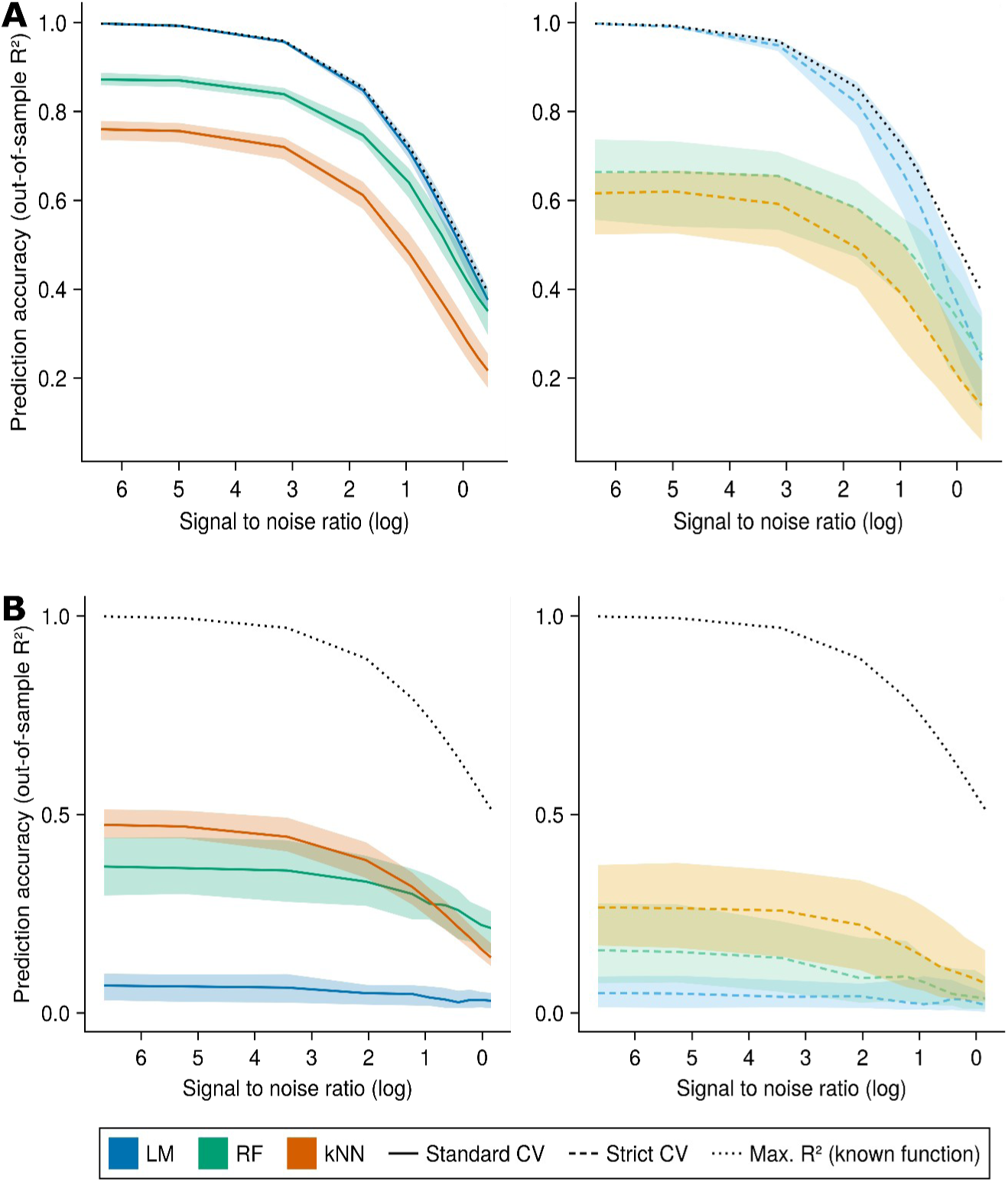
(A) Prediction accuracy over a range of signal-to-noise ratios for the three tested models under the assumption of a linear relationship between traits and yield. (B) Prediction accuracy over a range of signal-to-noise ratios for the three tested models under the assumption of a non-linear relationship between traits and yield.

Not unexpectedly, linear models failed to capture the more complex patterns driven by non-linear trait-overyielding relationships (Fig. 4B). Under standard cross-validation, kNN slightly outperformed random forests at high SNR, while RF outperformed kNN at lower SNR levels. However, both models again showed substantial drops in predictive ability between standard and strict cross-validation, indicating that data leakage is relevant also in a context where the true underlying relationship to be modelled is non-linear. Overall, our simulation suggest that the apparent model success in standard cross-validation is likely driven by component-level identity leakage.

To better understand the ecological relevance of these results, we compared the predictive performance of the models under linear and non-linear trait-yield relationships at a signal-to-noise ratio of around 1 (Fig. 5). When assuming a linear relationship, under standard cross-validation LM predicted 46.9% of the variation in overyielding, dropping to 34.9% under strict cross-validation. Similarly, the our-of-sample R^2^ of RF dropped from 41.9% to 32.1%, and that of kNN dropped from 28.2% to 19.3%. When simulating overyielding using the non-linear function, LM always performed poorly, with 3.3% of the variation in OY explained under standard cross-validation and 2.7% under strict cross-validation. The variation in OY explained by RF dropped from 22.2% to 4%, and the variation in OY explained by kNN dropped from 16.1% to 8.6% between standard and strict cross-validation.

**Figure 5:**
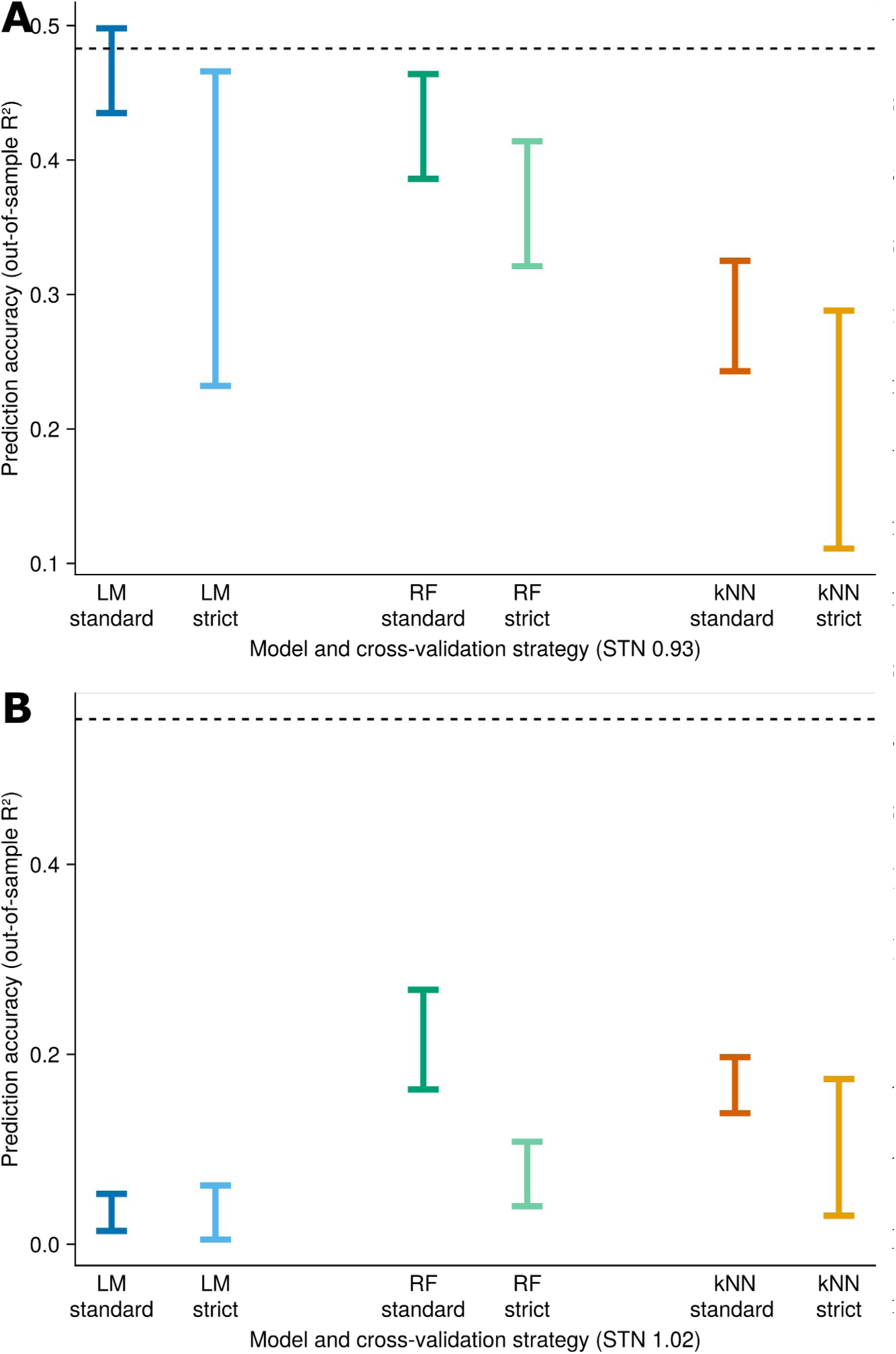
(A) Prediction accuracy for a signal-to-noise ratio of around 1 for the three tested models under the assumption of a linear relationship between traits and yield. (B) Prediction accuracy for a signal-to-noise ratio of around 1 for the three tested models under the assumption of a non-linear relationship between traits and yield.

The maximum R^2^ achievable by a correctly specified model, determined by fitting the same function used to generate the simulation data, was 48.3% for the linear relationship and 55.3% for the non-linear relationship. Our simulations show that achieving such R^2^ values in correct, i.e. strict cross-validation, is only possible when the signal is very strong. For example, for a SNR of around 2.5 and simulating a linear relationship between measured traits and yield, LM (using the best possible model after AIC-based backwards variable selection) under strict cross-validation achieved an out-of-sample R^2^ of 66.2%, while the maximum achievable R^2^ was 72.2%. Analogously, when the simulated relationship was non-linear, for a SNR of around 2.5 the best performing model, kNN, achieved 14.2%, while the maximum achievable R^2^ was 74.1%.

This emphasizes the need to distinguish between model fit (explained variance when fitting the model to the entire available data) and actual predictive power when assessing trait-based models of overyielding, and has direct implications for interpreting BEF studies that report high variance explained by (linear or non-linear) models without assessing generalizability through strict cross-validation.

## Discussion

Here, using soybean as a model system, we aimed to predict genotype mixture overyielding from traits determined in monoculture stands or on single plants. We compared two distinct cross-validation types to evaluate model performance. The first (“standard cross-validation”) considers mixtures that share a component as independent. The other cross-validation type we applied (“strict cross-validation”) tested predictions only on mixtures composed of genotypes different from those used in model training.

### Model performance depends on trait complexity and method flexibility

Even in relatively simple models (e.g., linear models using the best combination of predictors), the different cross-validation types can lead to substantially different interpretations of how well overyielding in new mixtures is predictable. Whereas the coefficient of determination of linear model fits or predictions based on standard cross-validation were in a range in accordance with previous reports (Roscher et al. 2012; Jochum et al. 2020; Montazeaud et al. 2020), the ones calculated from strict cross-validation were substantially lower.

For linear trait-overyielding relationships, linear models performed robustly and captured the main signal. However, when trait-function relationships were non-linear, non-parametric methods such as random forests or k-nearest neighbors can capture patterns that linear models miss, although they require more data and are more sensitive to noise (Table 1). Non-linear and interactive effects of functional traits on ecosystem productivity have been documented in several BEF studies (Holzwarth et al. 2015; Gonzalez et al. 2020), supporting the use of non-linear or non-parametric models to capture these complex relationships. This suggests that moderate predictive success of traits in previous studies (e.g. van der Plas et al. 2020) may partly reflect limitations of linear methods for capturing complex trait interactions rather than absence of a dependency on traits.

**Table 1:** Comparison oft the strengths and limitations of the three models compared when using them to predict mixture overyielding from mixture-components’ monoculture trait measurements.

| *Method* | *Strengths* | *Limitations* | *Data Requirements* | *Sensitivity to Leakage* | *Best use case* |
| --- | --- | --- | --- | --- | --- |
| *Linear* | Robust, easy to interpret, performs well with linear trait–overyielding relationships | Cannot capture non-linear interactions | Low | Low | Simple systems or known linear trait–function relationships |
| *Random Forest* | Flexible, captures complex interactions, handles non-linear effects | Requires large datasets to perform well | High | High | Complex trait interactions, large datasets, non-linear trait–function relationships |
| *kNN* | Flexible, captures local patterns, handles non-linear effects | Limited extrapolation, performs poorly with sparse data | High | High | Systems with high-quality data and strong non-linear patterns |

### Standard cross-validation overestimates predictive ability due to data leakage

The difference in performance between the two cross-validation methods can be explained by a special case of data leakage, appearing when there are “dependencies between training and test data that do not exist between training and inference-time data” (Bernett et al. 2024). Indeed, in standard cross-validation, the training and test data are to some degree correlated because some overyielding estimates, albeit in different mixtures in training and test datasets, are made relative to the same monoculture yield of a component genotype. Similarly, trait differences used as predictors in training models and during validation are calculated relative to the same measurement of a genotype trait. In strict cross-validation, on the other hand, the training and test data do not share any information at genotype level. Therefore, standard cross-validation is limited in its broader applicability as well as its suitability to inform fundamental ecological questions such as the relationship between functional trait differences and mixture productivity. More generally, this illustrates the distinction between explaining variation within a dataset versus predicting new, independent observations (Shmueli 2010): models may appear successful under standard cross-validation because they capture correlations within the observed data, but they can fail to generalize to novel genotype combinations. Strict cross-validation avoids this pitfall by providing a more realistic assessment of predictive power.

### Error propagation as an additional source of uncertainty

In addition to data leakage, the statistical propagation of measurement errors further reduces the signal-to-noise ratio in overyielding data, making prediction even harder. Because overyielding is derived from mixture performance and the average monoculture performance of its components, any measurement error in monocultures or mixtures propagates into the overyielding estimate. In our empirical data, we estimated that ∼46% of the variation in overyielding could potentially be explained if all measurements were independent. However, when accounting for the fact that many mixtures refer to the same monoculture measurements, this proportion dropped to ∼42%. While the difference may appear small, it highlights that shared measurement errors induce correlations among overyielding estimates of mixtures sharing monoculture references. This structural dependence is a form of error propagation that exacerbates leakage between training and test datasets, complicating model evaluation and prediction. As this issue is inherent as soon as multiple mixtures share monoculture references, experimentally it could only be avoided by growing unique monoculture references for each mixture, a design that is rarely feasible. Statistically the problem might be overcome by explicitly modelling the error correlation structures among the set of measurements, something that is typically not done. Lower signal-to-noise ratios caused by error propagation further exacerbate the challenges of developing predictive models and highlight the need for rigorous validation approaches. Together, these observations underscore the importance of rigorous validation and careful method selection for predictive modeling in ecological studies.

### Broader relevance and conclusions

Although our study focused on overyielding, the issue of predictive inflation due to data leakage is likely relevant across a broad range of ecological prediction tasks, from biomass production to ecosystem stability, wherever the units of prediction (e.g., mixtures, plots) share overlapping components or identities. Our simulations and empirical results show that predictive ability should be clearly distinguished from model fit, and validation protocols must avoid identity-based dependencies between training and test data.

Similar structural dependencies are common in ecological prediction tasks, where species or genotypes are repeatedly used across observations. Such non-independence can inflate apparent predictive success if not explicitly accounted for. Our simulations suggest that much of the apparent predictive success reported in prior studies may be inflated by identity-based data leakage rather than true generalization, particularly when noise levels are high.

At the same time, our results highlight that machine learning models such as random forests or k-nearest neighbors have considerable potential in BEF research, particularly when trait-yield relationships are complex and nonlinear, as is likely the case in many ecological settings. However, these models are also particularly sensitive to data leakage and can show highly inflated predictive performance under standard cross-validation (Yates et al. 2023). We therefore argue that adopting strict cross-validation as the standard will lead to more realistic assessments of both traditional linear models and machine learning approaches, helping to identify when apparent patterns are driven by leakage rather than genuine ecological signal. Altogether, this will ultimately improve the reliability of ecological predictions and the understanding of trait-function relationships in BEF research.

## Acknowledgments

This work was supported by the Swiss National Science Foundation, project 310030_192537. We acknowledge Jürgen Krauss and Enis Mathlouthi (both Agroscope) for technical support. We thank Claude-Alain Bétrix (Agroscope) and the USDA germplasm bank for providing seed material for our experiments. We acknowledge Christian Tancredi, Lydia Terrani, Sabrina Robbiani, Rahel Brühlmann, Timothy Dietrich and Jeremias Stalder for their help both on the field and in the greenhouse.

## Author contributions

E.B.K., N.K., S.E.W. and P.A.N conceptualized and designed the research. E.B.K, N.K., L.V. and S.E.W performed the experiments. E.B.K performed the simulations with input by P.A.N. E.B.K performed the analyses and wrote the manuscript with input from S.E.W and P.A.N. All authors revised and approved the final version of the manuscript.

---

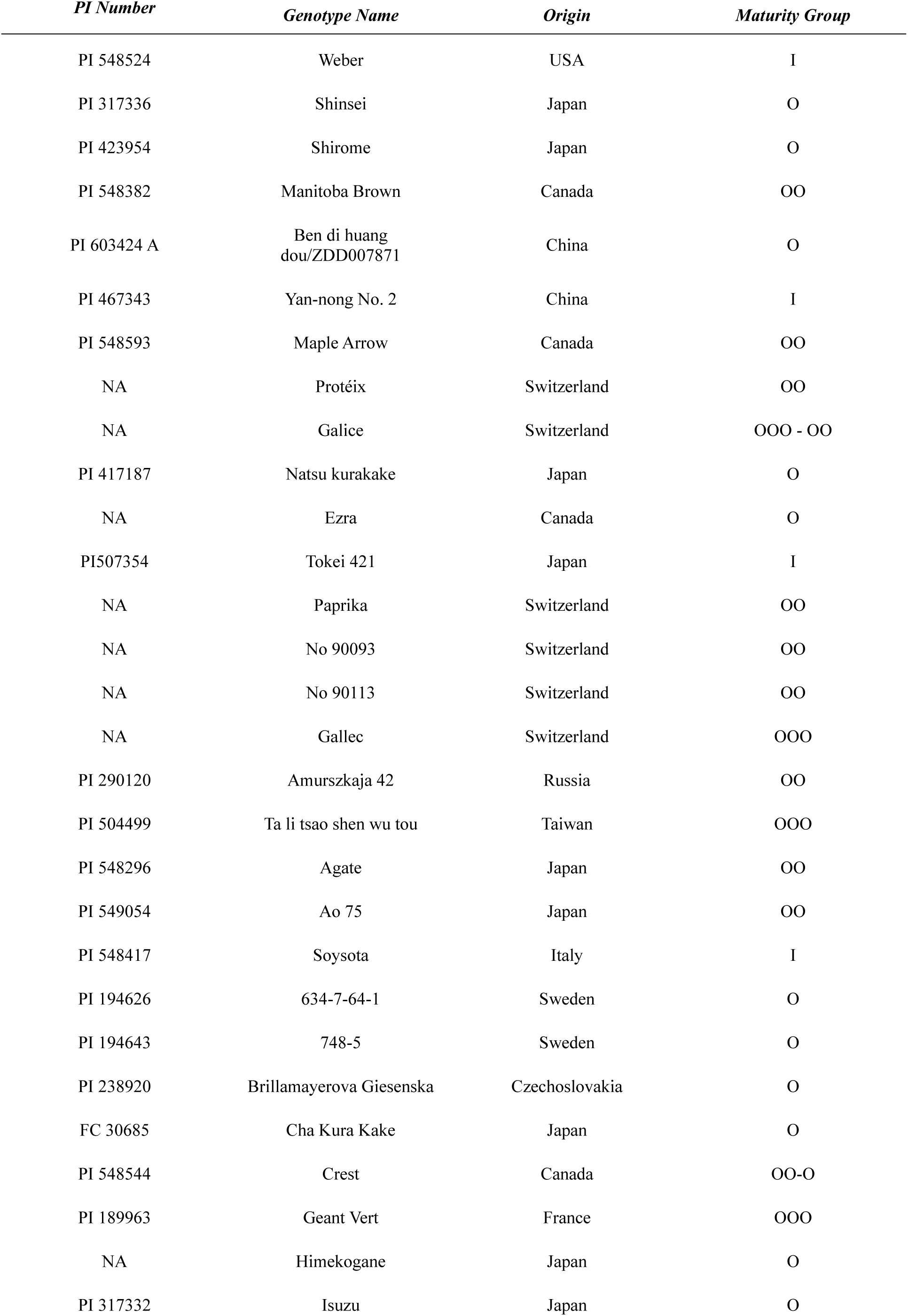
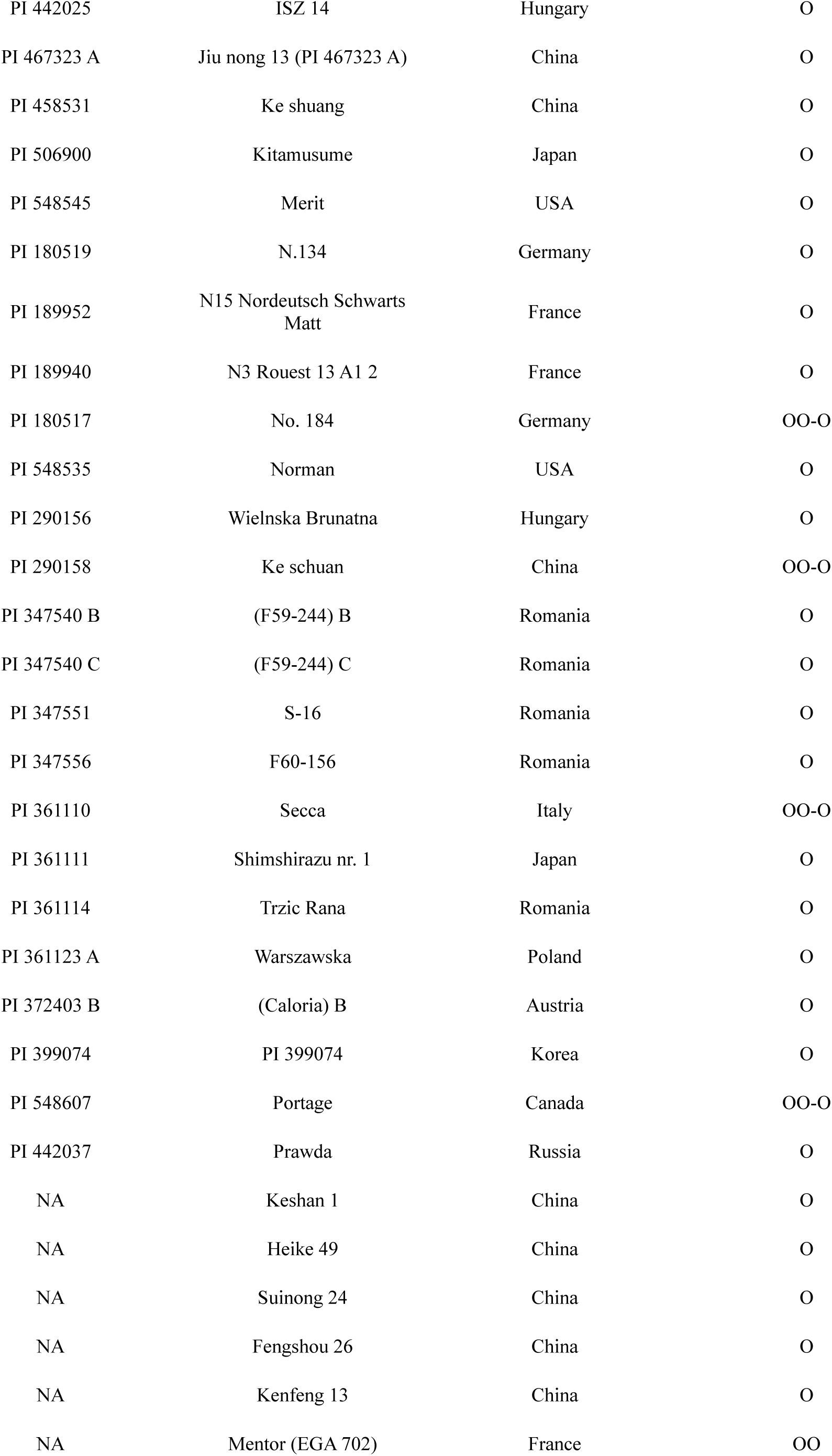
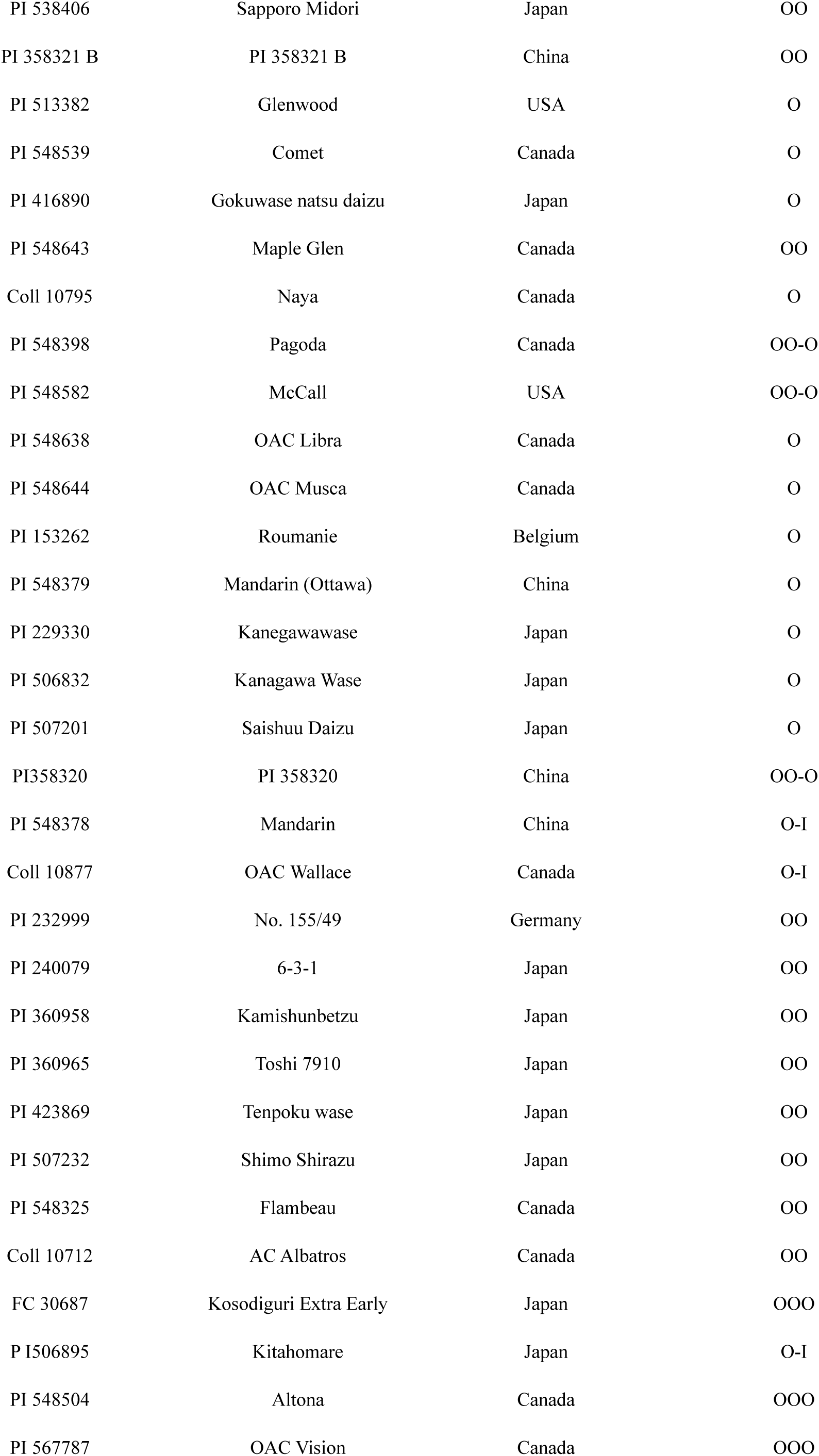

## References

Amyntas A, Berti E, Gauzens B, et al (2023) Niche complementarity among plants and animals can alter the biodiversity–ecosystem functioning relationship. Functional Ecology 37:2652–2665. 10.1111/1365-2435.14419

Barry KE, Mommer L, Ruijven J van, et al (2019) The Future of Complementarity: Disentangling Causes from Consequences. Trends in Ecology & Evolution 34:167–180. 10.1016/j.tree.2018.10.013

Bates D, Noack A, Kornblith S, et al (2023) JuliaStats/GLM.jl: v1.9.0

Bernett J, Blumenthal DB, Grimm DG, et al (2024) Guiding questions to avoid data leakage in biological machine learning applications. Nat Methods 21:1444–1453. 10.1038/s41592-024-02362-y

Blaom A, Lienart T, Simillides Y, et al (2024) alan-turing-institute/MLJ.jl: v0.20.3

Blonder B (2018) Hypervolume concepts in niche- and trait-based ecology. Ecography 41:1441–1455. 10.1111/ecog.03187

Borg J, Kiær LP, Lecarpentier C, et al (2018) Unfolding the potential of wheat cultivar mixtures: A meta-analysis perspective and identification of knowledge gaps. Field Crops Research 221:298–313. 10.1016/j.fcr.2017.09.006

Cardinale BJ, Wright JP, Cadotte MW, et al (2007) Impacts of plant diversity on biomass production increase through time because of species complementarity. Proc Natl Acad Sci USA 104:18123–18128. 10.1073/pnas.0709069104

Carlsson K, Karrasch D, Bauer N, et al (2024) KristofferC/NearestNeighbors.jl: v0.4.20

Chase JM, Leibold MA (2003) Ecological Niches: Linking Classical and Contemporary Approaches. University of Chicago Press

Ebeling A, Pompe S, Baade J, et al (2014) A trait-based experimental approach to understand the mechanisms underlying biodiversity–ecosystem functioning relationships. Basic and Applied Ecology 15:229–240. 10.1016/j.baae.2014.02.003

Fixt E, Hodges JL (1989) Discriminatory Analysis. Nonparametric Discrimination: Consistency Properties

Godoy O, Gómez-Aparicio L, Matías L, et al (2020) An excess of niche differences maximizes ecosystem functioning. Nat Commun 11:4180. 10.1038/s41467-020-17960-5

Gonzalez A, Germain RM, Srivastava DS, et al (2020) Scaling-up biodiversity-ecosystem functioning research. Ecology Letters 23:757–776. 10.1111/ele.13456

Hector A, Schmid B, Beierkuhnlein C, et al (1999) Plant Diversity and Productivity Experiments in European Grasslands. Science 286:1123–1127. 10.1126/science.286.5442.1123

Ho TK (1995) Random decision forests. In: Proceedings of 3rd international conference on document analysis and recognition. IEEE, pp 278–282

Holzwarth F, Rüger N, Wirth C (2015) Taking a closer look: disentangling effects of functional diversity on ecosystem functions with a trait-based model across hierarchy and time. R Soc open sci 2:140541. 10.1098/rsos.140541

Hooper DU, Chapin FS, Ewel JJ, et al (2005) EFFECTS OF BIODIVERSITY ON ECOSYSTEM FUNCTIONING: A CONSENSUS OF CURRENT KNOWLEDGE. Ecological Monographs 75:3–35. 10.1890/04-0922

Huang T, Döring TF, Zhao X, et al (2024) Cultivar mixtures increase crop yields and temporal yield stability globally. A meta-analysis. Agron Sustain Dev 44:28. 10.1007/s13593-024-00964-6

Huang Y, Chen Y, Castro-Izaguirre N, et al (2018) Impacts of species richness on productivity in a large-scale subtropical forest experiment. Science 362:80–83. 10.1126/science.aat6405

Hutchinson GE (1957) Concluding Remarks. Cold Spring Harbor Symposia on Quantitative Biology 22:415–427. 10.1101/SQB.1957.022.01.039

Jochum M, Fischer M, Isbell F, et al (2020) The results of biodiversity–ecosystem functioning experiments are realistic. Nat Ecol Evol 4:1485–1494. 10.1038/s41559-020-1280-9

Kopp EB, Niklaus PA, Wuest SE (2023) Ecological principles to guide the development of crop variety mixtures. Journal of Plant Ecology 16:rtad017. 10.1093/jpe/rtad017

Mahaut L, Violle C, Shihan A, et al (2023) Beyond trait distances: Functional distinctiveness captures the outcome of plant competition. Functional Ecology 37:2399–2412. 10.1111/1365-2435.14397

Montazeaud G, Violle C, Roumet P, et al (2020) Multifaceted functional diversity for multifaceted crop yield: Towards ecological assembly rules for varietal mixtures. J Appl Ecol 57:2285–2295. 10.1111/1365-2664.13735

Reiss ER, Drinkwater LE (2018) Cultivar mixtures: a meta-analysis of the effect of intraspecific diversity on crop yield. Ecol Appl 28:62–77. 10.1002/eap.1629

Roberts DR, Bahn V, Ciuti S, et al (2017) Cross-validation strategies for data with temporal, spatial, hierarchical, or phylogenetic structure. Ecography 40:913–929. 10.1111/ecog.02881

Roscher C, Schumacher J, Gubsch M, et al (2012) Using Plant Functional Traits to Explain Diversity–Productivity Relationships. PLoS ONE 7:e36760. 10.1371/journal.pone.0036760

Sadeghi B, Poom Chiarawongse, Squire K, et al (2022) DecisionTree.jl - A Julia implementation of the CART Decision Tree and Random Forest algorithms

Shmueli G (2010) To Explain or to Predict? Statist Sci 25:. 10.1214/10-STS330

van der Plas F, Schröder-Georgi T, Weigelt A, et al (2020) Plant traits alone are poor predictors of ecosystem properties and long-term ecosystem functioning. Nat Ecol Evol 4:1602–1611. 10.1038/s41559-020-01316-9

Violle C, Jiang L (2009) Towards a trait-based quantification of species niche. Journal of Plant Ecology 2:87–93. 10.1093/jpe/rtp007

Wagg C, Ebeling A, Roscher C, et al (2017) Functional trait dissimilarity drives both species complementarity and competitive disparity. Funct Ecol 31:2320–2329. 10.1111/1365-2435.12945

Wuest SE, Peter R, Niklaus PA (2021) Ecological and evolutionary approaches to improving crop variety mixtures. Nat Ecol Evol 5:1068–1077. 10.1038/s41559-021-01497-x

Yates LA, Aandahl Z, Richards SA, Brook BW (2023) Cross validation for model selection: A review with examples from ecology. Ecological Monographs 93:e1557. 10.1002/ecm.1557

